# A Systematic Identification of RBPs Driving Aberrant Splicing in Cancer

**DOI:** 10.1101/2023.07.17.549307

**Authors:** Marian Gimeno, César Lobato-Fernández, Ane San Martín, Ana Anorbe, Angel Rubio, Juan A. Ferrer-Bonsoms

**Author notes:** Correspondence: Juan A. Ferrer-Bonsoms, Angel Rubio. These authors contributed equally and share first authorship (Marian Gimeno, César Lobato-Fernández). These authors share senior and last authorship (Angel Rubio, Juan A. Ferrer-Bonsoms).

## Abstract

Alternative Splicing (AS) is a post-transcriptional process by which a single RNA can lead to different mRNA and, in some cases, several proteins. Various processes (probably many of them yet to be discovered) are involved in the regulation of alternative splicing. This work focuses on the regulation by RNA-binding proteins (RBPs). In addition to splicing regulation, these proteins are related to cancer prognosis and are emerging therapeutic targets for cancer treatment. CLIP-seq experiments target selected RBPs and result in uncovering the loci of the nascent transcriptome to where the RBP binds to. The presence of changes in the splicing status surrounding these loci is a good starting point to establishing a causal relationship. The selection of the specific RBP(s) to target in the CLIP-seq experiment is not straightforward; in many cases, this selection is driven by *apriori* hypotheses.

In this work, we have developed an algorithm to detect RBPs that are likely related to the splicing changes between conditions. To do this we have integrated several databases of CLIP-seq experiments with an algorithm that detects differential splicing events to discover RBPs that are especially enriched in these events. This is a follow-up of a previous work that is refined by 1) improving the algorithm to predict the splicing events and 2) testing different enrichment statistics, and 3) performing additional validation experiments. As a result, the new method provides more accurate predictions, and it is also included in the Bioconductor package EventPointer.

We tested the algorithm in four different experiments where seven different RBPs were knocked down. The algorithm accurately states the statistical significance of these RBPs using only the alterations in splicing. We also applied the algorithm to study sixteen cancer types from The Cancer Genome Atlas (TCGA). We found relationships between RBPs and several cancer types like *CREBBP* and *MBNL2* alterations in adenocarcinomas of the lung, liver, prostate, rectum, stomach, and colon cancer. Some of these relationships have been validated in the literature but other ones are novel.

**Availability:** This method is integrated EventPointer, an available Bioconductor R package.

## 1 Introduction

Splicing is a co-transcriptional process that removes introns and splices the exons to generate mature mRNA from pre-mRNA(1). In turn, if there is Alternative Splicing (AS), the patterns of exons retained in the mature mRNA (and therefore, the introns removed) change under different conditions. These different mRNA species are called isoforms or transcripts and, in turn, can lead to different proteins. These proteins may also have different functions. All the hallmarks of cancer (angiogenesis, immortality, immune response avoidance, etc.) have been related to aberrant alterations in AS (1).

AS regulation is a complicated process that involves the spliceosome. The spliceosome is a complex composed of small nuclear RNAs, proteins, and polypeptides assembled jointly with the transcription process. The proteins that interact with RNA are called RNA-binding proteins (RBPs). Most proteins in the spliceosome are also RBPs and more than 1500 RBPs (2) can be identified in the human species. These proteins regulate many processes related to RNA such as RNA metabolism, RNA methylation (m6A (3)), and, of course, alternative splicing. Cross-linking and immunoprecipitation sequencing (CLIP-seq) experiments are the method of choice for experimental validation of the actual interaction of an RBP with the mRNA. These experiments require knowing in advance which RBP must be targeted.

RBPs are known to play an important role in cancers. Mutations or changes in their expression can influence oncogenes’ expression levels. In fact, RBPs can be targeted when developing cancer treatments (4). Hence, it would be very useful to know which are the RBPs related to changes in the splicing (5).

In a previous work (6), we developed an algorithm that, by integrating different databases of CLIP experiments and differential splicing analysis, infers the potential RBPs conducting these changes in splicing. The underlying procedure identifies the RBPs that bind to genomic loci near the differential splicing events and states which RBPs are significantly enriched. This algorithm only requires RNA-seq information to predict the RBPs involved in the splicing changes. The accurate selection of RBP candidates that drive the splicing requires selecting the database that includes the CLIP-seq experiments, using a proper enrichment statistical method, an algorithm to detect the AS, and also fine-tuning different hyperparameters.

In this work, we have upgraded the previous method in all these aspects. We have included the recently published POSTAR3 database (7) (which includes 32% more RBP experiments than the previous version for human and mouse). We have selected a more accurate algorithm to detect the differential splicing events, and included different statistical methods –Hypergeometric, GSEA, Wilcoxon, and Poisson-Binomial– to state the enrichment of the RBPs. We validated this algorithm on real data where some RBPs were knocked down and consistently provided better results than the previous algorithm.

Further, we applied it to 19 different types of cancer on the TCGA and TARGET database to predict cancer-specific RBPs. This analysis shows several relationships between RBPs and cancers, some of them are known –for instance, *CREBBP* and *MBNL2* alterations in adenocarcinomas of the lung and liver respectively (8,9)– and others are novel. In addition to including it in Bioconductor, we also developed a Shiny application to ease the analysis of the results and extract conclusions for the scientific community.

## 2 Results

The main result is the implementation of the algorithm itself: SFpointer is a resource that pinpoints RNA-Binding Proteins (RBPs) especially enriched in zones where there is differential splicing.

This objective is achieved by integrating several CLIP-seq databases into a single one to perform enrichment analysis on the RBPs. The result is a ranked list of RPBPs that are potentially driving the changes in the splicing pattern of an experiment.

The development of this software involved several challenges: i) Since CLIP-seq is not as widely used as RNA-seq, the prediction of SFs binding motifs strongly depends on the availability of CLIP experiments, so that, an extensive integration of databases is desirable, ii) the statistical significance depends both in the quality of the call of splicing events and the statistical method used in the enrichment analysis, iii) The results of the statistical pipeline must be validated on experiments where the ground truth is known. The algorithm must be easy to use and available to the scientific community. We have successfully resolved these challenges.

### 2.1 Included RBPs are increased by 30% with the updated databases

Compared to the previous version, we have included POSTAR3 (7), a recently published large CLIP-seq database (**Figure 1.1**). POSTAR3 contains 1445 CLIP experiments from 7 different species covering 348 RBPs. We have downloaded and curated only experiments mapped against human and mouse genomes due to the similarity of these species. Results from the mouse genomes were mapped to the human genome (using liftover (10)) to get their equivalent in the human genome(version GRCHg38). Total number of included RBPs –from human and mouse– is 244, i.e. 25% of the number of searchable RBPs in the previous version (6).

**Figure 1:**
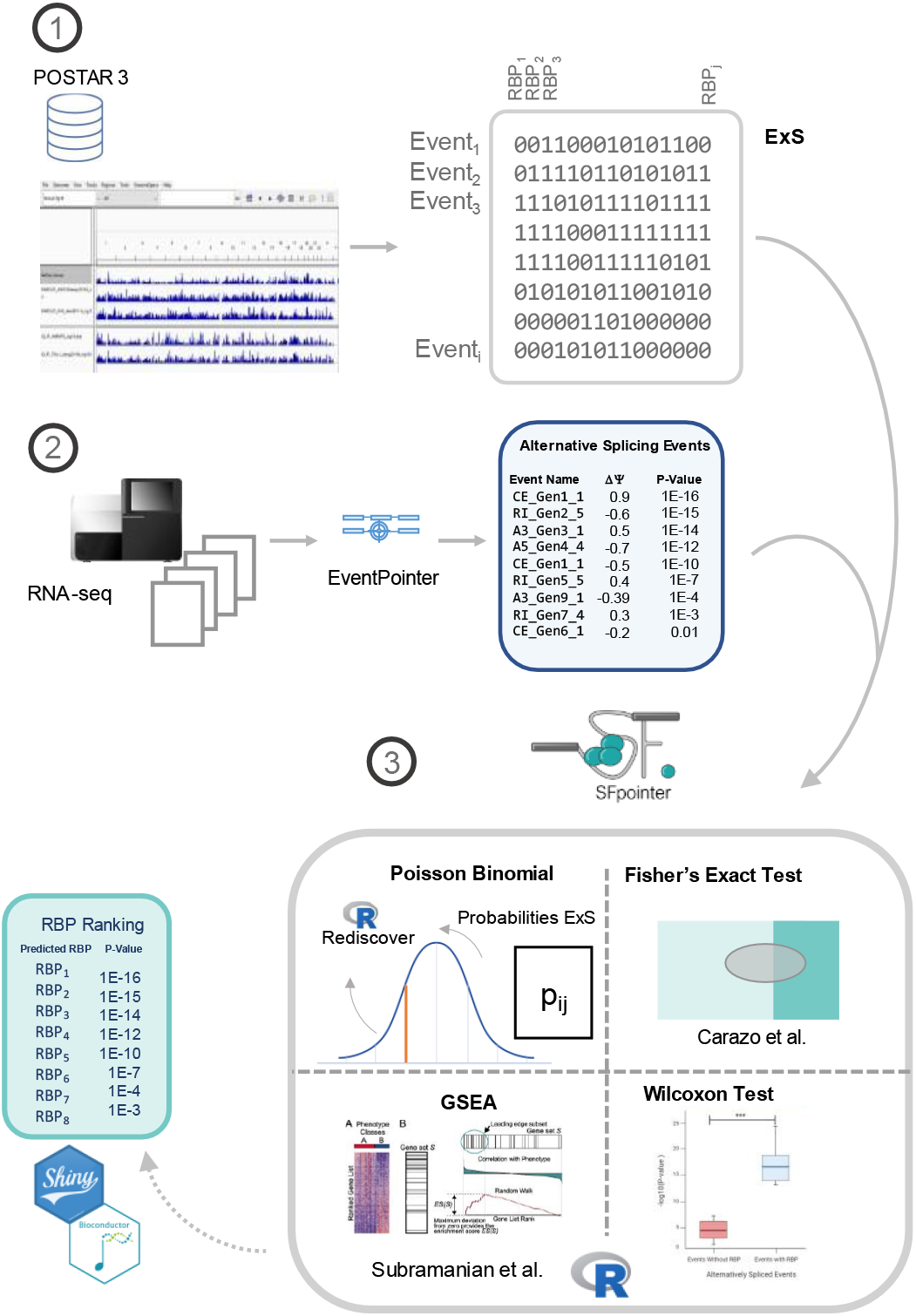
SFpointer Pipeline. **1) Building the *ExS* matrix.** CLIP experiments from POSTAR 3 were translated into ***ExS***, which is a matrix whose entry *i, j* is 1 if the genomic loci where the RBP “j” binds, is in the neighborhood of the splicing event “i” in any of the CLIP experiments included in the database, and a 0 otherwise. **2) Alternative Splicing Study**. The differentially spliced events are detected using a bootstrap version of EventPointer. **3) Context-specific RBPs**. To estimate the enrichment of the RBPs, SFpointer receives as input the differentially spliced events and the ***ExS*** matrix. It performs one of the four available enrichment methods: Poisson Binomial, Fisher’s Exact Test, GSEA, and Wilcoxon Test, returning as output a ranking of RBPs with the corresponding enrichment p-value. This method has been implemented in a shiny APP and Bioconductor.

This integrated database provides genomic loci that were mapped to the corresponding genome (hg38 in our case). In EventPointer (11), the potential splicing events were annotated based on a reference transcriptome (in this one GENCODE24). We compared the RBP binding sites and the potential splicing events. To establish the relationship between an RBP binding site and a splicing event, we created an indicial sparse matrix denoted as ***ExS*** (Events x Splicing Factors). This matrix states whether a splicing event has an RBP binding locus in its neighborhood (within a 400 nt window of the event). The exact details of the ***ExS*** construction can be consulted in our previous work (6).

### 2.2 Boosting sensitivity and specificity through a new statistical modeling

Regarding the computation of the input data, we noticed that the correct identification of the altered splicing events strongly impacts the performance of the SFs calculation. A proper selection of the differential splicing events makes the enrichment of RBPs more precise. We have switched to a bootstrap-based statistic recently implemented in the EventPointer package (**Figure 1.2**). This statistical approach increases the sensitivity compared to the previous version (11).

**Figure 2:**
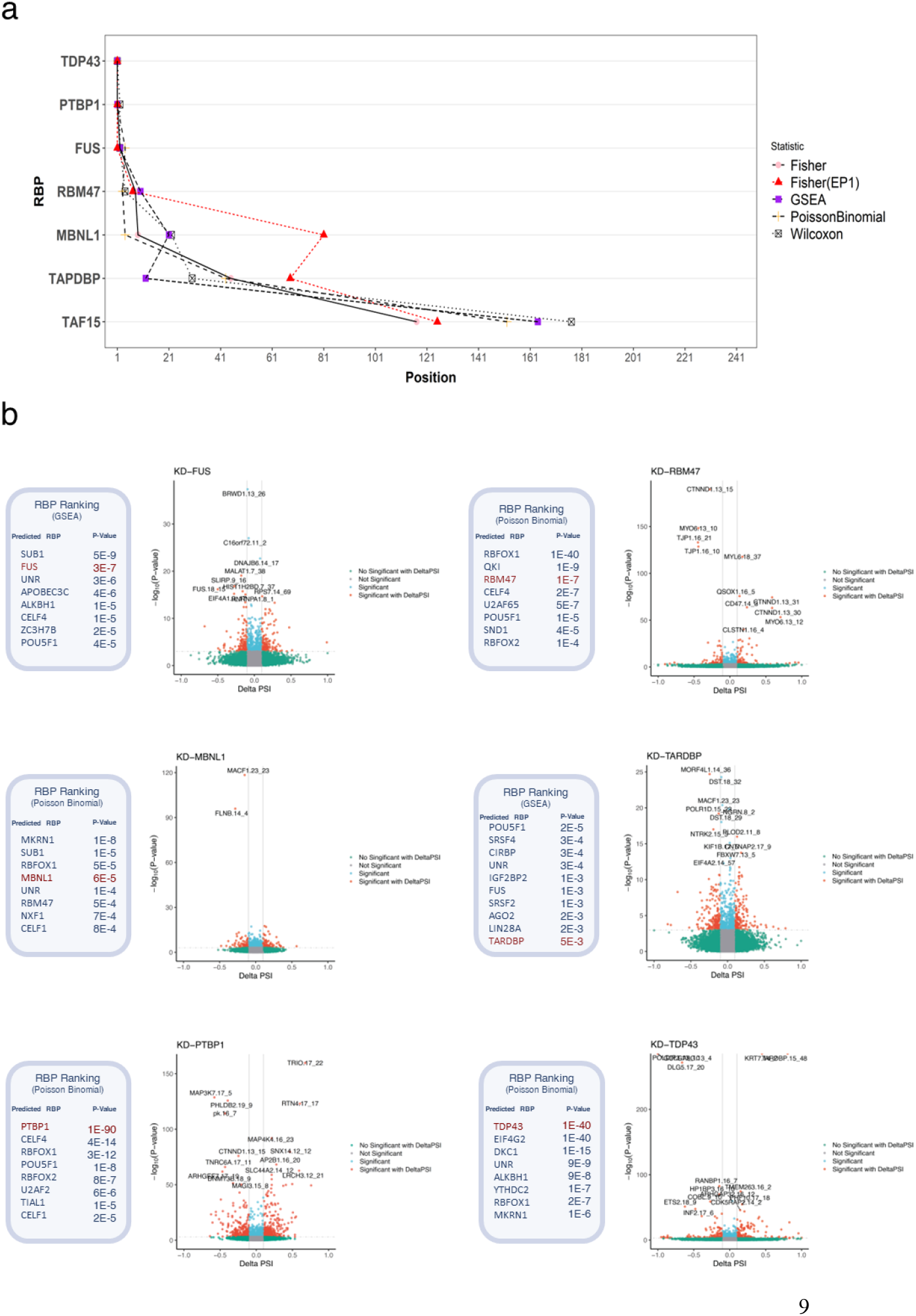
SFpointer validation across 7 independent KD experiments. **a) Positions of the RBPs for each of their KD using the original version and four methods included in SFpointer.** Fisher, GSEA, Poisson Binomial, and Willcoxon are shown in pink, purple, black, and orange respectively, and the original version of SFpointer with the previous pipeline of EventPointer but using the current ***ExS*** is shown in red. Each point represents the ranking position of each RBP for the different statistical approaches. The KD-*TAF15* experiment is included as evidence of the absence of alternative spicing activity. **b) Individual results for each of the conditions**, excluding KD-*TAF15* due to its low effect on alternative splicing. For each condition, the top 8 RBPs (top 10 in KD-*TARDBP*) and their enrichment p-values are included. The enrichment method shown is the one that improves the RBP’s position in the ranking. A volcano plot with the results of the AS analysis for each condition is included along with the ranking of RBPs. This plot shows in red color those AS events that have an absolute value of Delta PSI (ΔΨ) greater than 0.1 and a P-value less than 1e-3, in blue color, are shown those significant events without such a noticeable change in DeltaPSI and in green color are shown those events with a large change in Delta PSI but which are not significant.

Secondly, we implemented four different statistical enrichment analyses on the AS events to predict the differential activity of the RBPs (**Figure 1.3**). The first method is based on the hypergeometric distribution using a Fisher’s Exact Test. This test estimates, using the ***ExS*** matrix, the enrichment of the splicing factors (SFs) by setting a threshold on the p-values to state which are the differentially spliced events. This method is consistently used to perform GO enrichment analysis and was already implemented in the previous version of the algorithm (6,12,13).

**Figure 3:**
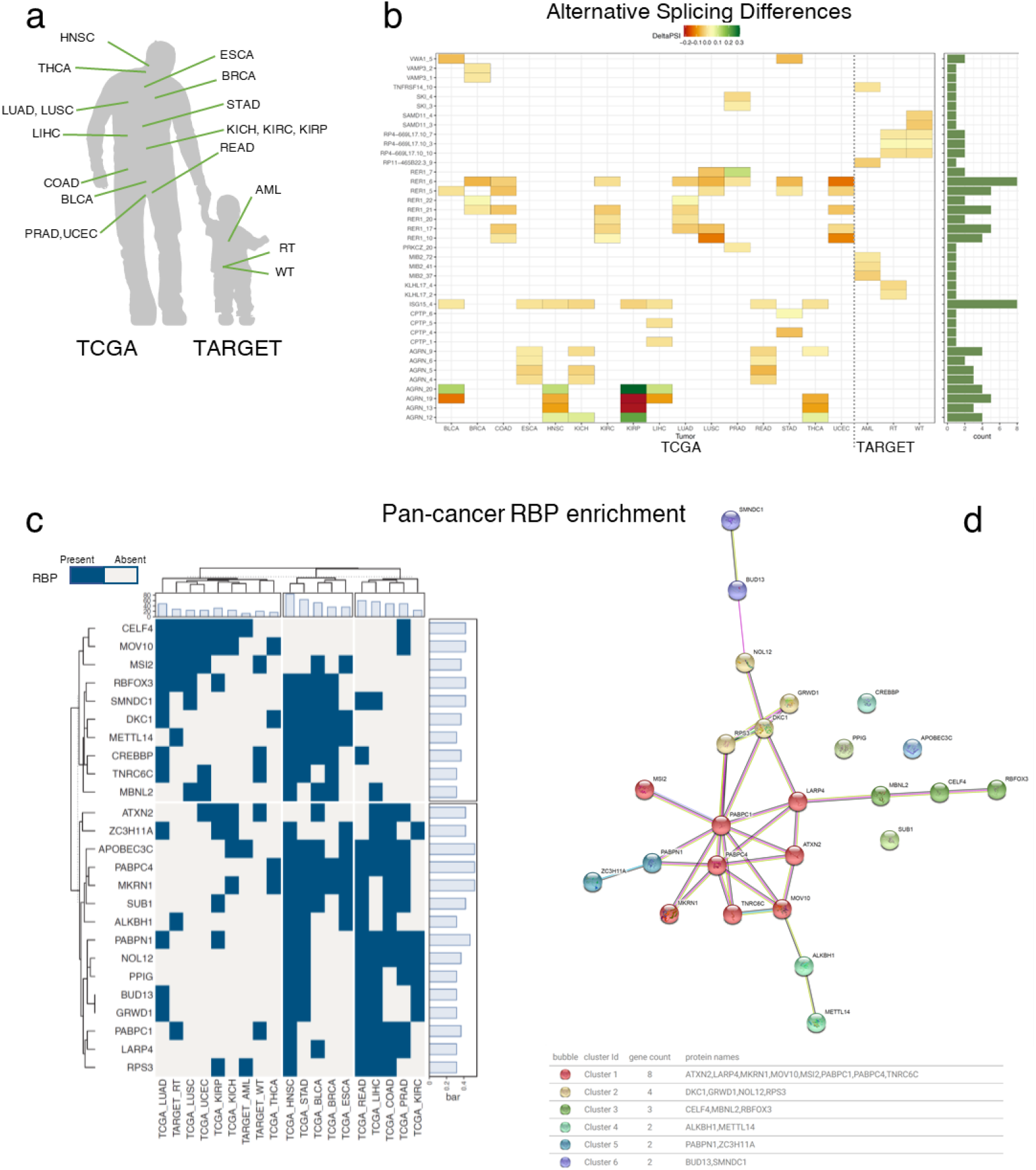
Pan-Cancer Study of Splicing Regulators. **A) Figure showing the tumor types included in the pan-cancer study.** These tumor types belong to TCGA and TARGET. Only those tumor types with enough normal samples to perform the AS study have been included. **B) Pan-cancer AS analysis**. Heatmap showing the differences in alternative splicing. This graph shows on the abscissa axis the different cancer types and on the ordinate axis each of the splicing events. These events are selected by taking the top 5 most significant splice events for each cancer site. The figure shows the ΔΨ for each event in each condition, the red color is a negative ΔΨ and the green color a positive delta PSI. **C) Pan-cancer RBP enrichment**. Heatmap showing the 22 RBPs significantly enriched in more than 5 tumor types. These RBPs have been clustered into 2 groups and in blue color are shown the tumors in which they appear enriched. These cancer sites appear in three different clusters. A row and a column bar chart showing the abundance of RBP in different tumor types and the number of RBPs per tumor type is shown. **D) STRING Clustering**. This graph shows the relationship of RBPs according to the STRING database. The colors refer to each of the clusters, each of the bubbles represents an RBP and the lines joining them are the evidence of the relationship provided by STRING. The table at the bottom of the graph includes the description of these clusters.

The data used to perform the enrichment analysis (the matrix ***ExS***) shows some potential sources of bias: some RBPs bind to a large proportion of the splicing events (and other RBPs scarcely bind to any splicing event). Furthermore, there are some splicing events with many hits from RBPs and other ones with very few. A similar statistical analysis was performed in Discover (14) for the detection of mutually exclusive mutations. In that study, it was shown that the differences in the density of both the rows and the columns of the input matrix induce a bias in a naïve analysis based on the hypergeometric distribution.

In order to correct for this bias, we used Rediscover(15), an R package that implements the Poison-Binomial distribution instead of a plain hypergeometric distribution. Both the hypergeometric and the Poison-Binomial approaches, require the users to select a threshold to state when a p-value is considered significant.

None of the two previous methods exploits the ranking of the aberrant AS events: a splicing event that appears in the 1^st^ position is equally important –from the point of view of the previous algorithms– as one that appears in the last position as long as the p-value is below the threshold. GSEA(16), an enrichment analysis based on the Kolmogorov-Smirnov test, exploits this information and it was shown to have a very good statistical power in GO enrichment analysis. As a result, it is also included in the package. Finally, we have also included a standard Wilcoxon test, a non-parametric test that also exploits the ranking of the events.

### 2.3 SFpointer accurately identifies the RBP causing splicing disruption in 7 KD experiments

We tested the proposed pipeline on 4 RNA-seq experiments with 7 different knocked-down RBPs, allowing us to test its accuracy (**Figure 2**). We rerun these experiments using the previous version of SFpointer to compare the statistical approach. To make a fair comparison, we used the current ***ExS*** matrix on the previous version of the algorithm (**Table 1**). This way, it is possible to identify if the changes in the approach (both the enrichment method and the selection of splicing events) improved the overall sensitivity and precision of the current method.

**Table 1:** Table showing the ranking positions for the 7 KD-RBP conditions. The results compare the positions in the ranking using the original version with the current ExS and the four new enrichment methods with the current EventPointer pipeline and ExS. The minimum positions in ranking for each condition are highlighted in bold.

|  | *RBP* | *SFpointer*<br>*Original*<br>(Including<br>*POSTAR3*) | *SFpointer*<br>*New*<br>(Fisher's<br>Exact Test) | *SFpointer*<br>*New*<br>(Poisson<br>Binomial) | *SFpointer*<br>*New*<br>(GSEA) | *SFpointer*<br>*New*<br>(Wilcoxon<br>Test) |
| --- | --- | --- | --- | --- | --- | --- |
| *PRJEB39343* | PTBP1 | **1** | **1** | **1** | **1** | 2 |
| *PRJEB39343* | MBNL1 | 81 | 9 | **4** | 21 | 22 |
| *GSE77702* | FUS | **1** | 2 | 4 | 2 | 2 |
| *GSE77702* | TAF15 | 125 | **117** | 152 | 164 | 177 |
| *GSE77702* | TARDBP | 68 | 45 | 43 | **10** | 30 |
| *GSE136366* | TDP43 | **1** | **1** | **1** | **1** | **1** |
| *GSE75491* | RBM47 | **7** | 8 | **3** | 10 | 4 |

We have restudied the analysis included in (6), which tested the ability of the algorithm to predict two different SFs from GSE77702. In this dataset, we compared three different contrasts: KD-*FUS*, KD-*TARDBP*, and KD-*TAF15* vs scramble transfection (17). Results in the previous version -using POSTAR 2-predicted *FUS* the 10^th^ ranked in KD-*FUS* and *TARDBP* the 33^rd^ ranked in KD-*TARDBP*(6). Using this new approach, FUS is ranked 2^nd^ for KD-*FUS* (8 positions above the original study) and *TARDBP* was ranked the 10^th^ for KD-*TARDBP*. Only, by updating the ***ExS*** matrix, results with the original version ranked 1^st^ for KD-*FUS* and 68^th^ for KD-*TARDBP*. Results in the KD-*TARDBP* condition might seem concerning, but expression analysis revealed that *TARDBP* knockdown failed. The KD procedure did reduce the levels of gene expression, but it did not knock-down completely the gene (**Figure S2**). Finally, KD-*TAF15* was excluded from the previous study due to the low impact that the activity of this RBP has on alternative splicing, previously demonstrated in other articles (6,17). This is in agreement with the results detected with our pipeline in which KD-*TAF15* appears below position 100 out of 244 for any of the statistical approaches (**Figure 2a**).

To further validate the approach, we applied this methodology to three additional datasets. In the first one, (*PRJEB39343*) in which three RBPs (*PTBP1, ESRP2*, and *MBNL1*) were knocked-down in gastric cancer lines (18). We performed our approach excluding the KD-*ESRP2* condition because it is not contained in the actual ***ExS***. Focusing the analysis on KD-*PTBP1* and KD-*MBNL1*, our pipeline precisely pinpointed *PTBP1* to be the most likely altered RBP and *MBNL1* ranked in the top 5 in their respective condition that differ from the original version of the algorithm which pinpointed *PTBP1* but left *MBNL1* to the 81^st^ position (**Figure 2b**;**Table 1**).

The second and third experiments include KD-*TDP43* (GSE136366)(19) and KD-*RBM47* (GSE75491) (20) conditions, performed on HeLa and H358 cell lines respectively. For these two additional conditions, SFpointer successfully ranks *TDP43(TARDBP) as* 1^st^ and ranks *RBM47* in the 3^rd^ position versus the original version that ranked 1^st^ and 7^th^ respectively. Note that *TARBP* and *TDP43* are the same gene with very different ranking results, which reinforces the statement that the enrichment results are strongly dependent on the quality of the experiment. In *GSE136366* the expression of *TARDBP* decreases 10 times while in experiment *PRJEB39343* it only decreases 2 times (Figures S2 and S6).

**Figure 2a** includes a comparison of the enrichment methods, for which the position in the ranking of the targeted RBP is shown. The figure shows that the new AS calculation pipeline for EventPointer (Fisher) increases the sensitivity versus the previous approach (Fisher’s Exact Test EP1). Furthermore, the new enrichment methodologies GSEA, Wilcoxon, or Poisson Binomial also increases the precision over the Fisher’s approach. As could be noticed, results depend strongly on the experiment as it could be easily comprehended, that when there is a deficiency in AS alteration, the RBP is not achieving a high score in the ranking (KD-*TAF15*, KD-*TARDBP*). Nevertheless, the other RBPs ranked in the top 5 positions with strongly significant p-values (**Figure 2b**).

All of the conditions were ranked in the top 10 out of 244 positions proving successfully the detection power. In addition, all the enrichment methods provide good results. The Poisson Binomial distribution is marginally better (**Table 1**). Full results from the enrichment analyses are included in **Supplementary Tables 1-6**.

Finally, for each KD experiment, **Figure 2b** includes an AS analysis result that showed the AS alteration on each condition and is possibly related to the RBP splicing regulator activity. For instance, *BRWD1* or *MALAT-1* are lncRNA with a strongly significant decrement in Ψ for KD-*FUS* condition, which has been previously reported in relation to *FUS* activity (21). Also, *FLNB* in KD-*MBNL1* has been described as part of the pathway of *MBNL1*-mediated apoptosis (22), and *MAPK* kinases genes in KD-*PTBP1*condtion which has been reported as a *MARPK/ERK* pathway inhibitor (23).

### 2.4 Pan-cancer analysis of splicing regulators reveals three groups of tumors with similar RBPs profiles

It has been proved in several studies the relevance of aberrant splicing in cancer development(12,24,25). Here we have included a pan-cancer alternative splicing (AS) study conducted to identify RBP dis-regulation similarities across 19 different cancer types and 9,514 patients. We analyzed all TCGA and TARGET tumor samples containing normal samples (**Figure 3a**).

As a starting point we studied the aberrant AS in each cancer type using the EventPointer pipeline –and bootstrap statistics– and once we obtained those splicing events with a distinct pattern, we ran for each condition an enrichment in RBPs using the four enrichment tests that SFpointer incorporates. The results shown in **Figure 3** are those obtained from the enrichment using only the Poisson Binomial approach. We finally clustered the most frequent RBPs using STRING (26).

**Figure 3b** shows the top 5 alternatively spliced events of each cancer type from TCGA and TARGET. It can be noticed that several AS events are recurrent in different types of cancer and that AS events in childhood-driven cancer are very specific for each cancer type. Consequently, there are no co-occurrent AS events shared between these diseases and other AS adult cancers. On the other hand, two main genes in the top 5 positions for each cancer type are recurrently differentially spliced across adult tumors: *AGRN* (ENSG00000188157) and *RER1* (ENSG00000157916). AS events concerning *AGRN* appear in 4 out of 16 adult cancers: ESCA, KICH, READ, and THCA. *AGRN* is a gene whose isoform expression is tissue-specific and it has been recently linked to the Hippo pathway in the tumor microenvironment for different cancer sites (27,28), and its aberrant splicing is related to impaired neuromuscular junction synaptogenesis (29) although there are no current studies linking its splicing with tumorigenesis.

In the case of *RER1*, AS events in this gene ranked top 5 in 9 out of 16 tumor types, with event number six having a special relevance in 8 out of the 16 TCGA types. Interestingly, the *RER1* gene has been associated with colon and pancreatic cancer (30,31). More specifically, one of its AS events has been reported in association with disease relapse in colon cancer (31) and its biological function has been reported for carcinogenesis induction in pancreatic cancer(30).

Furthermore, using the results obtained by SFpointer on each cancer site, we selected the RBPs that appeared significantly enriched in at least 5 different cancer sites. We performed a k-means clustering by RBPs and cancer sites with 10-fold cross-validation. The results are included in **Figure 3c**. This Figure shows 3 different clusters regarding the cancer types according to the amount of RBPs dis-regulated in each condition being: HNSC, STAD, BLCA, BRCA, and ESCA the tumor types with the highest number of altered RBPs, having all of them in common the alteration of *DKC1, METTL14, PABPC4*, and *MKRN1*. All of them have a strong relationship with cancer development, i.e. *DKC1* is related to the expression of tumor suppressors (32), *METTL14* mediates tumor progression through *SOX4* alteration and *WTAP* (3), *PABPC4* is downregulated in metastatic cells (33), and *MKRN1* modulates tumor progression by AKT signaling pathway (34).

The second cluster includes relevant cancer sites such as COAD or READ and shares the alteration of *PABPN1*, and *NOL12*, both of them related to tumor progression (35–37). Finally, the third group includes the tumors with the fewest dis-regulated RBPs in which LUAD and TARGET cancer sites are also included. This group is characterized by the high presence of altered *CELF4* and *MOV10* among its samples. Both genes have been related to cancer development (38,39).

Besides considering the clustering by RBPs functionality in pan-cancer, there are two main clusters i) one that is leading AS alterations in adult cancer (bottom cluster in the plot), this group includes relevant cancer genes e.g. *MKRN1, DKC1*, or *PABPC4*; and ii) the other one is modulating AS in more tissue-specific sites (top part of the plot) and comprises relevant oncogenes e.g. *CREBBP* (8) or *MBNL2* (9).

Finally, with the list of the 22 RBPs that were selected for the previous analysis, we clustered the RBPs using STRING MCL methodology (26), finding 6 clusters shown in **Figure 3d** and in **Supplementary Table 7**, e.g. cluster 1 comprises *LARP4, ATXN2, MKRN1, PABPC1, PABPC4, MOV10, TNRC6C*, and *MSI2* RBPs; and cluster 2 contains *DKC1, NOL12, GRWD1*, and *RPS3* RBPs. Besides, comparing the clusters obtained using STRING annotation and the results obtained by our method, we see that of the 19 RBPs clustered by STRING, 13 have been included in the same group (roughly 70%).

### 2.5 Constructing a pan-cancer splicing regulators resource

We implemented the results obtained for the pan-cancer RBP enrichment and the AS analysis results in a developed shiny app that can be accessed through (https://gitlab.com/Jferrerb/sfpointer_gui). The shiny app enables the cancer site selection and contains the complete ranking with the 244 RBPs for 19 different tumor types extracted from TCGA and TARGET.

This app allows viewing the results of alternative splicing analysis for each of the 16 conditions shown in **Figure 2a**. In addition, it is possible to visualize each of the specific events.

Enrichment results are also included for each of the RBPs, which can be queried at the RBP level (see in which tumors it appears as enriched) and by condition (see all the RBPs enriched in a tumor type). Moreover, it is possible to download the graphs and tables from the app itself and perform a survival analysis according to each RBP in each tumor type, to get an overview of the relevance of each RBP in its contribution to the overall survival.

Furthermore, the code has been integrated into the EventPointer package already available in the Bioconductor repository. The code vignettes and a model of the pipeline are included in https://github.com/clobatofern/SFPointer_testPipeline.git.

## Discussion

In this study, we have developed and implemented a new method to detect potential RBPs drivers at AS in different biological conditions. Results can be directly inferred from an RNA-seq experiment allowing to calculate the disruption of 244 RBPs –avoiding the need for performing 244 CLIP experiments. Furthermore, SFpointer has been validated using 7 different KD experiments. Results of the validation presented the disrupted RBP in the top 5 of predicted ones and outperformed the previous methodology. Finally, we have applied it to TCGA and TARGET discovering pan-cancer actuation RBPs and new cancer-specific RBPs that have been made available for consultation by any user through our Shiny app.

Our software is a statistical method that is based on co-occurrence, but we are aware that it does not imply causality. We cannot claim, using the plain results, that the predicted RBPs are causing the observed splicing changes. It is a method that only states that the genomic loci where some particular RBPs bind, are especially enriched in places where there is differential splicing. Henceforth, it provides an educated guess to perform some type of biological validation of the involved RBPs.

In reference to the above, the method relies for its predictions on the ***ExS*** matrix that relates the genomic loci of the RBPs with the alternative splicing events of the transcriptome. This matrix was constructed using all the human and mouse experiments from POSTAR3. It includes CLIP experiments from many different conditions and tissues were stored in the database. However, apart from translation to GRCh.38, no further normalization was performed. Thus, SFpointer predicts over a particular tissue experiment using information from cell lines of other tissues, which is debatable since each tissue has a very different behavior. However, we have prioritized predictive ability over prediction accuracy, i.e. for a certain condition we prefer to have the possibility to give a result than to reduce the predictive ability to 1 or 2 RBPs, because of the scarcity of CLIP experiments performed on those cell lines. In six out of the seven experiments, this approach proved to be valid.

Finally, regarding enrichment methods, we have included in our tool most of the state-of-the-art methods: Fisher’s Exact Test, Poisson Binomial, GSEA, and Wilcoxon. The first two methods do not consider the rankings of events with alternative splicing, while the latter does. As expected, the results of the four methods are quite similar, and all of them perform reasonably well. We noticed that the precision of the RBP prediction strongly depends on the conditions of the experiment -i.e. *TARDBP* is predicted in 10^th^ and 1^st^ position in two different KD-*TARDBP* experiments being the first a less effective KD of *TARDBP*- and in these conditions, the enrichment methods tend to differ in the results obtained. The robustness of the AS analysis will also considerably affect the result, we recommend the use of EventPointer as its results are robust and we have been able to validate them, e.g. by identifying the MEK pathway with *PTBP11*.

About the validations using 7 KD experiments, we applied each of the four methods and the results demonstrate that the statistical advances presented in this work improve the results obtained with the previous version of SFpointer. Indeed, we observe that in the experiments with no technical failures, the enrichment results place the RBP in the top 5 of the ranking of alerted RBPs. Remarkably, the four enrichment methods provide similar predictions in each condition. Although GSEA enrichment and Poisson Binomial especially stand out for their performances, the former is an order of magnitude slower, but both are equally accurate and have obtained the best qualitative result of the validations.

A major contribution of this article is the application of the SFpointer pipeline to all data from both TCGA and TARGET. The results obtained are very promising: we have achieved the identification of two co-occurring splicing events present across different tumor types. This fact highlights the relevance of the study of splicing concerning cancer, and how it could be possible to include splicing events as biomarkers (40,41). Isoform-specific data was downloaded from (42) that used as reference GENCODE24. This is the reason why we used a somehow old version of the transcriptome.

Likewise, we have performed an enrichment in RBPs for the different tumor types, obtaining three different groups of behavior depending on the number of RBPs in which they are enriched, having a special variability of splicing in tumors such as BRCA, HNSC, while pediatric tumors or lung cancer have much less variability in the enrichment of RBPs. The presence of *PABPC4* and *MKRN1* has been observed as the most frequently enriched RBPs in the different types of cancer coherently with literature (33,34), proving the relevance of this approach.

Besides, approximately 70% of the RBPs predicted with our methodology cluster similarly using STRING data and analytics. Interestingly, using a completely different information, we have deduced a qualitatively similar behavior. This gives a glimpse of the statistical power of the method.

Finally, we have developed a shiny application through which it is possible to consult the results of the pan-cancer analysis, the events with which a binding site of an RBP coincides, and the RBPs that have a binding site in a given AS event. This app is available at (https://gitlab.com/Jferrerb/sfpointer_gui). We have also added code and the corresponding vignettes with their explanation to Bioconductor, where it is integrated within EventPointer for use by all those researchers who wish to give a first biological interpretation of the results of their alternative splicing analysis.

## 4 Methods

### 4.1 Data availability

RBPs binding information was extracted from a recently upgraded resource POSTAR3(7), publicly available at (http://postar.ncrnalab.org).

Fastq files corresponding to the GSE136366 and the GSE75491 experiments were downloaded from ENA. Transcript expression was obtained from the Fastq files using Kallisto. Alternative splicing analysis was performed using the EventPointer pipeline. TCGA data were downloaded from (42), where transcript expression was computed using Kallisto and GENCODE24 as reference transcriptome.

The Code of all the analyses is available in (https://github.com/clobatofern/SFPointer_testPipeline.git). Results of regarding the TCGA data can be consulted in the previously mentioned shiny app (https://gitlab.com/Jferrerb/sfpointer_gui).

### 4.2 Relationship between RBPs and Splicing Events

In this work, we started with our previously published work on Splicing Factor (SF) prediction (6). We collected 937 CLIP-Seq experiments for 244 different RBPs contained in POSTAR3(7). The building of the **ExS** matrix is identical to the one described in our previous work (6). We performed a change in genome version using the liftOver R package (10) to transform different genome versions from human and mouse species e.g. Hg19, mm9, mm10, into a human genome hg38. With this process, we increased the sample size for the human species.

After obtaining all binding sites in hg38, we mapped the binding sites against all the splicing events from transcriptome GeneCode v.24 calculated using the EventPointer pipeline (11). We stored this information in a sparse matrix denoted **ExS** (Events x Splicing factors). Each element denotes whether the splicing factor *j* binds to the event *i* as follows:

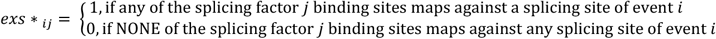

We have added some changes to improve the performance of the algorithm and integrated it into a Bioconductor R package. This version enables us to use the Fisher’s Exact Test, GSEA, a Wilcoxon test, and a new approach developed by us: the Poisson Binomial Enrichment.

### 4.3 Event statistics

Using **ExS** it is possible to perform an enrichment analysis on the differentially spliced events. We implemented EventPointer 3.0 bootstrap statistics for alternative splicing events detector. The main strength of the pipeline is the fact that it estimates the Ψ distribution for each event using bootstrap resulting in a very robust pipeline.

### 4.4 Methodology for RBP Enrichment and Ranking

Our method outputs a ranking with the most likely enriched RBPs.This ranking is performed using four different enrichment methods. We describe these methods in the following paragraphs (15).

#### 4.4.1 Fisher’s Exact Test

Fisher’s Exact Test was already described in (6). The Fisher test is based on a hypergeometric variable to calculate the probability of seeing an abnormal number of events that are differentially spliced and bound to an RBP. (**Equation 1**)

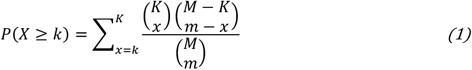

where,

- **M** is the total number of events
- **K** is the number of selected events.
- **m** is the number of events regulated by RBPi
- **k** is the number of events within K regulated by RBPi

We include in the current work two different options to select the relevant splicing events: select the splicing events with a Local False Discovery Rate under a threshold (Local FDR_thrs_=0.1) or select the first 1000 splicing events ranked by *p*-value. We used the first option in our pipeline.

#### 4.4.2 Poisson Binomial

From **ExS** matrix we compute the probability of a specific event being regulated by a specific RBP. The event-RBP regulation probability was estimated using the methodology proposed by (15). This methodology demonstrates that, assuming the independence between events and RBPs, the probability of an event i being regulated by an RBP j (P*ij*) can be written as 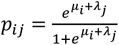 (for more details see (15)). Then, this approach uses a logistic regression as depicted in **Equation 2**:

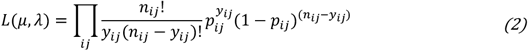

Where:

- **p_ij_** is the probability of event *i* being regulated by RBP *j*.
- P_**ij**_ can be written as 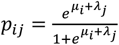 as depicted above.
- **n**_**ij**_ is the total number of cases: by construction is equal to 1.
- **y**_**ij**_ is equal to one if event *i* is regulated by RBP *j*, i.e., y_ij_ = {0,1}.

Then, we calculate the probability of observing an anormal number of events that are differentially spliced and regulated by an RBP. This probability is computed with the Poisson Binomial Distribution (**Equation 3**):

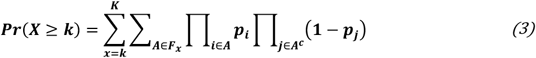

Where,

- **F**_**x**_ is the subset of *x* integers possible. if total number of elements is 3 and *x=2* then F_2_={{1,2},{1,3},{2,3}}
- **p**_**i**_ is the probability of an event *i* being regulated by a Splicing Factor.
- **P**_**j**_ is the probability of an event *j* being regulated by a Splicing Factor.

This is solved using Rediscover (15), which uses the poibin R package (43) to compute the p-values.

#### 4.4.3 Gene Set Enrichment Analysis (GSEA)

GSEA is a successful enrichment analysis method initially described in (16). GSEA is a non-parametric test based on the Kolmogorov-Smirnov statistic that compares the distributions of a variable (usually a p-value, but other possibilities are also valid) between the analytes (usually genes) that have a characteristic (usually a GO annotation) and those that do not have the characteristic. The application to RBP analysis is straightforward. The variable is the p-value of the splicing event and the characteristic is the presence or absence of an annotated RBP binding site in the neighborhood of the event.

We have used the R-package fgsea. It allows to “quickly and accurately calculate arbitrarily low GSEA P-values for a collection of gene sets” (44) by using an adaptive multi-level split Monte-Carlo scheme. Despite being the fastest available, it is still slower than any of the other implemented methods. One of the advantages of GSEA is that it does not require setting a threshold on the p-value to state which are the significant events.

#### 4.4.4 Wilcoxon’s Test

The Wilcoxon test can also be applied to perform an enrichment analysis. A Wilcoxon test is a non-parametric test that compares the medians of two datasets. In this case, the distributions to be compared are the p-values of the event annotated with an RBP binding site with the p-values of the events not annotated with the same RBP binding site.

The Wilcoxon test does not either require setting a threshold on the p-values. Our implementation (which uses sparse matrices and linear algebra) is the fastest of all the methods.

### 4.5 TCGA and TARGET analysis

We run the SFpointer pipeline using EventPointer 3.0 and the four enrichment analyses. For the alternative splicing analysis, we selected the top 5 differentially spliced events for each condition and extracted the delta PSI for each event to plot the comparative analysis.

Regarding the SFenrichment analysis, we extracted the RBPs that were present in at least 5 different cancer conditions and clustered them using Kmeans with 10-fold validation in two different groups. Additionally, we clustered the different cancer sites into 3 different groups using Kmeans and 10-fold cross-validation provided by the R package ComplexHeatmap (45).

Finally, with the obtained 22 RBPs present in more than 5 cancer conditions we used STRING MCL approach to plot and cluster the network, using STRING database information (44). The MCL inflation parameter was set to 3.

## Abbreviations

AML: Acute Myeloid Leukemia
BLCA: Bladder Urothelial Carcinoma
BRCA: Breast invasive carcinoma
COAD: Colon adenocarcinoma
ESCA: Esophageal carcinoma
HNSC: Head and Neck squamous cell carcinoma
KICH: Kidney Chromophobe
KIRC: Kidney renal clear cell carcinoma
KIRP: Kidney renal papillary cell carcinoma
LIHC: Liver hepatocellular carcinoma
LUAD: Lung adenocarcinoma
LUSC: Lung squamous cell carcinoma
PRAD: Prostate adenocarcinoma
READ: Rectum adenocarcinoma
RT: Rhabdoid Tumor
STAD: Stomach adenocarcinoma
THCA: Thyroid carcinoma
UCEC: Uterine Corpus Endometrial Carcinoma
WT: Wilms Tumor

## 5 Conflict of Interest Statement

The authors declare that the research was conducted in the absence of any commercial or financial relationships that could be construed as a potential conflict of interest.

## 6 Author contributions

**c**

## 7 Funding

The authors gratefully acknowledge the funding of the Editor project (Cancer Research UK [C355/A26819], AECC, and AIRC under the Accelerator Award Programme), PIBA_2020_1_0055 (funded by the Basque Government), and Synlethal Project (RETOS Investigacion, Spanish Government).

## 8 Acknowledgments

The results published here are in part based upon data generated by the TCGA Research Network: https://www.cancer.gov/tcga and Therapeutically Applicable Research to Generate Effective Treatments (https://ocg.cancer.gov/programs/target) initiative, phs000218.

The authors acknowledge the authors of POSTAR3 for providing the data from their database. The authors wanted to acknowledge Katyna Sada for helping with the SFpointer logo design.

## 9 Supplementary Material

Supplementary materials and tables are in an external file.

## Notes

https://gitlab.com/Jferrerb/sfpointer_gui

### Competing Interest Statement

The authors have declared no competing interest.

