## Supplementary_Material for "A Systematic Identification of RBPs Driving Aberrant Splicing in Cancer"

Supplementary Material: A Systematic Identification of RBPs Driving Aberrant Splicing in Cancer

†These authors share authorship

SUPPLEMENTARY MATERIAL

### 1 Supplementary Figures of KD expression

#### 1.1 GSE77702

The following pictures show the expression of FUS, TARDBP, and TAF15:

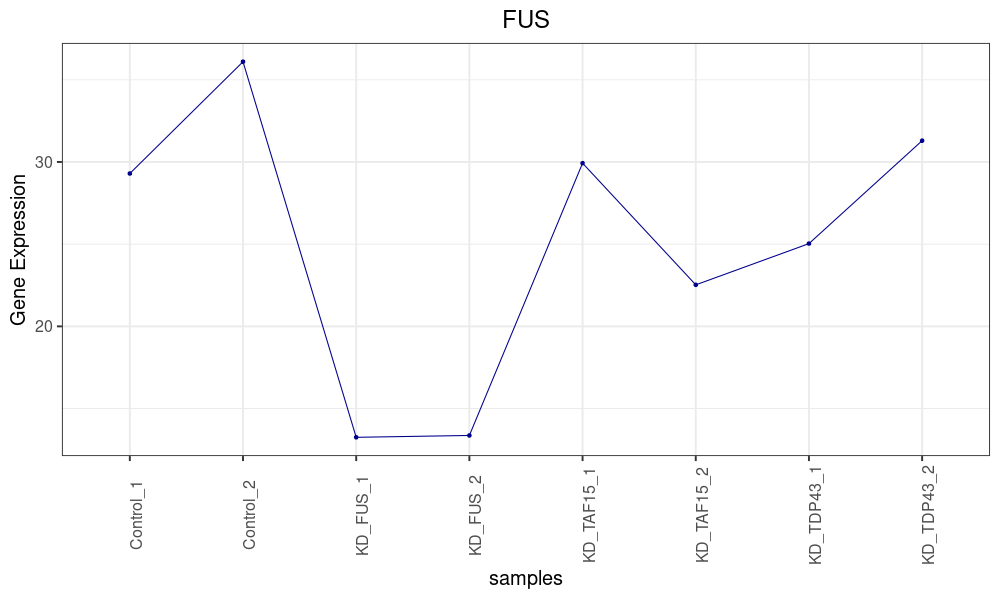

Figure S1. Expression of the FUS gene throughout the samples of the experiment GSE77702. The second and third samples correspond to the samples in which FUS was knocked down.

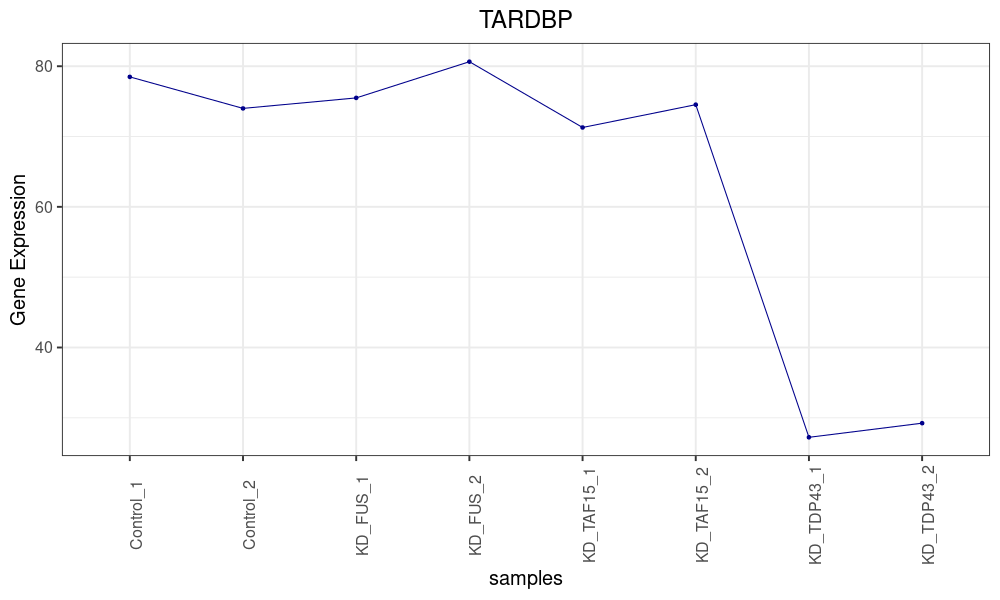

Figure S2. Expression of the TARDBP gene throughout the samples of the experiment GSE77702. The seventh and eighth samples correspond to the samples in which TARDBP was knocked down.

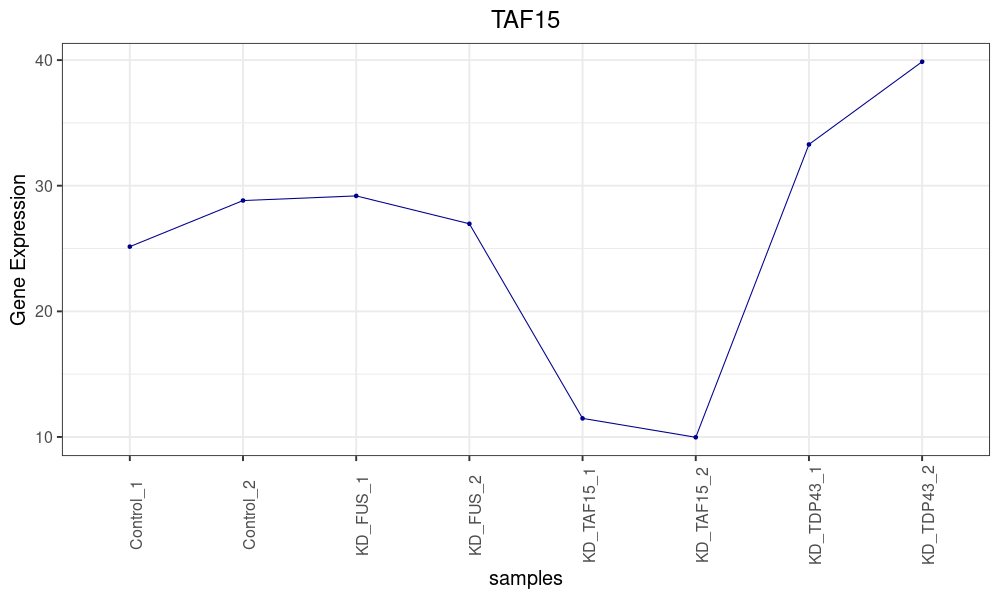

Figure S3. Expression of the TAF15 gene throughout the samples of the experiment GSE77702. The Fifth and sixth samples correspond to the samples in which TAF15 was knocked down.

**PRJEB39343**

The following pictures show the expression of MBNL1, and PTBP1:

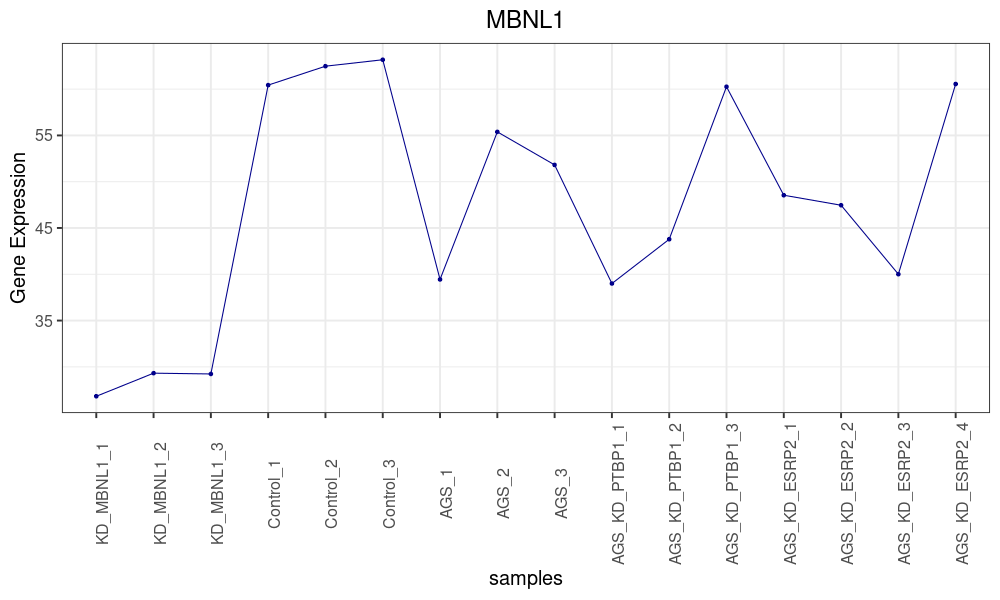

Figure S4. Expression of the MBNL1 gene throughout the samples of the experiment PRJEB39343. The first three samples correspond to the samples in which MBNL1 was knocked down.

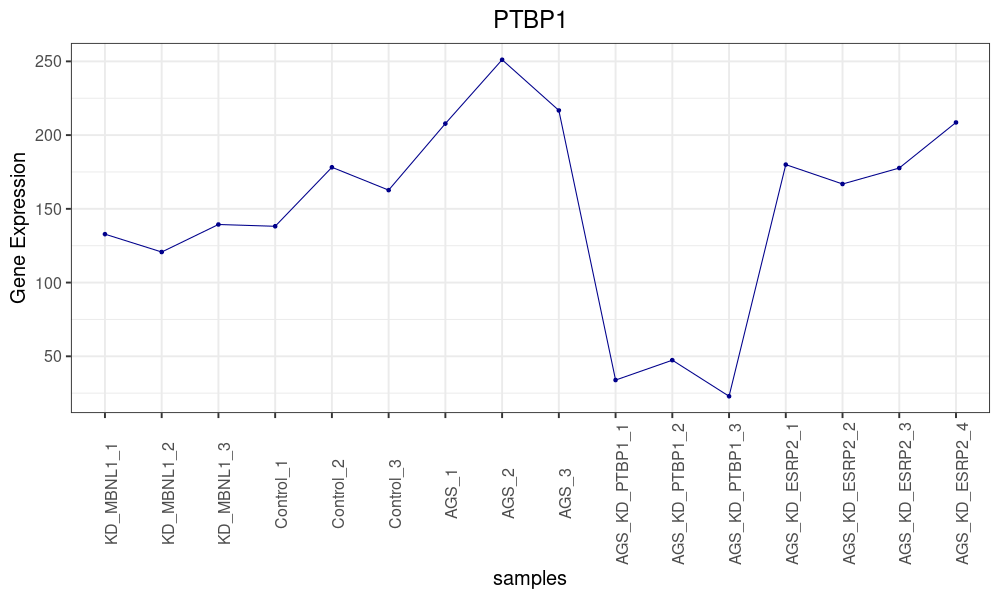

Figure S5. Expression of the PTBP1 gene throughout the samples of the experiment PRJEB39343. The tenth, the eleventh, and the twelfth samples correspond to the samples in which PTBP1 was knocked down.

**GSE136366**

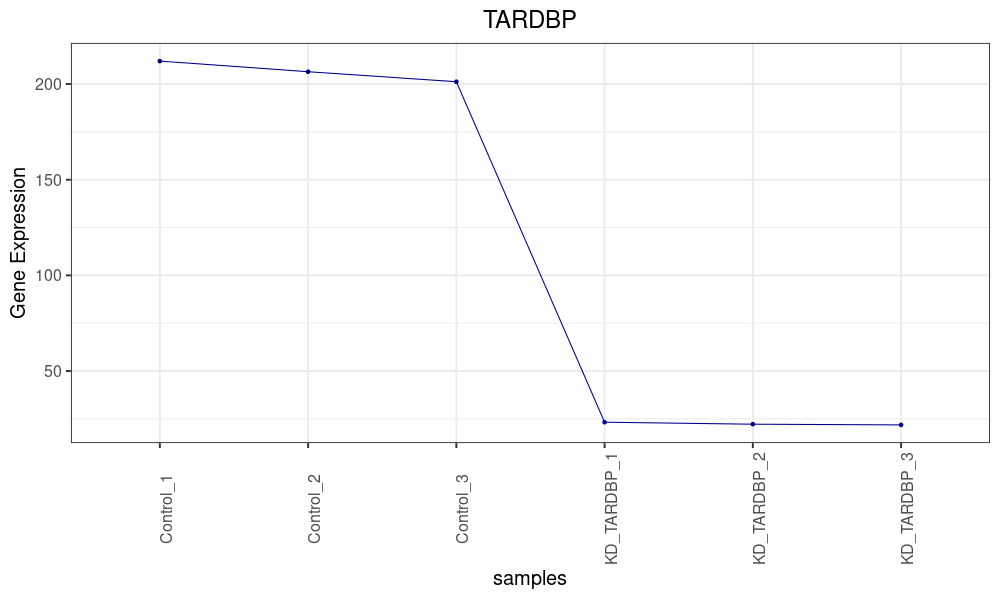

Figure S6. Expression of the TARDBP gene throughout the samples of the experiment GSE136366. The last three samples correspond to the samples in which TARDBP was knocked down.

**GSE75491**

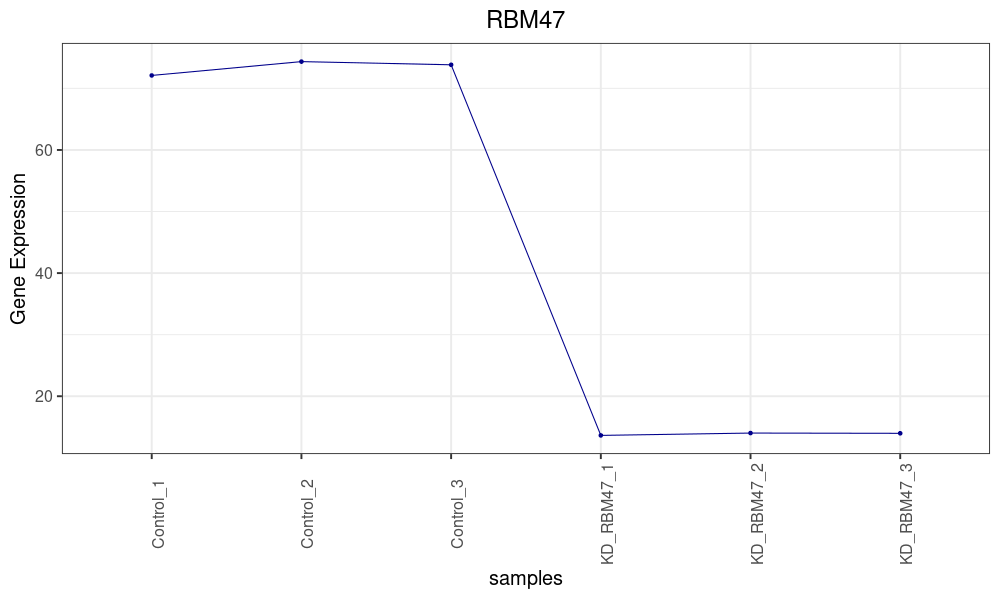

Figure S7. Expression of the RBM47 gene throughout the samples of the experiment GSE75491. The last three samples correspond to the samples in which RBM47was knocked down.
